# OGT binds a conserved C-terminal domain of TET1 to regulate TET1 activity and function in development

**DOI:** 10.1101/125419

**Authors:** Joel Hrit, Cheng Li, Elizabeth Allene Martin, Mary Goll, Barbara Panning

## Abstract

Mammalian TET enzymes oxidize 5-methylcytosine to 5-hydroxymethylcytosine and higher oxidized derivatives. TETs are targets of the enzyme OGT, which post-translationally modifies intracellular proteins in response to cellular nutrient status. The biological implications of the OGT-TET interaction have not been thoroughly explored. Here, we show for the first time that modification of TET1 by OGT enhances its activity *in vitro*. We identify a previously uncharacterized domain of TET1 responsible for binding to OGT and report a point mutation that disrupts the OGT-TET1 interaction. Finally, we show that the interaction between TET1 and OGT is necessary for TET1 to rescue *tet* mutant zebrafish hematopoetic stem cell formation, suggesting that OGT promotes TET1’s function in development. Our results demonstrate regulation of TET activity by OGT *in vitro* and *in vivo*. These results link metabolism and epigenetic control, which may be relevant to the developmental and disease processes regulated by these two enzymes.

## Introduction

Methylation at the 5’ position of cytosine in DNA is a widespread epigenetic regulator of gene expression. Proper deposition and removal of this mark is indispensable for normal vertebrate development, and misregulation of DNA methylation is a common feature in many diseases [1,2]. The discovery of the Ten-Eleven Translocation (TET) family of enzymes, which iteratively oxidize 5-methylcytosine (5mC) to 5-hydroxymethylcytosine (5hmC), 5-formylcytosine (5fC), and 5-carboxylcytosine (5caC), has expanded the epigenome [3-7]. These modified cytosines have multiple roles, functioning both as transient intermediates in an active DNA demethylation pathway [6,8-11] and as stable epigenetic marks [12,13] that may recruit specific readers [14].

One interesting interaction partner of TET proteins is *O*-linked N-acetylglucosamine (*O*-GlcNAc) Transferase (OGT). OGT is the sole enzyme responsible for attaching a GlcNAc sugar to serine and threonine residues of over 1,000 nuclear, cytoplasmic, and mitochondrial proteins [15,16]. Like phosphorylation, *O*-GlcNAcylation is a reversible modification that affects the function of target proteins. OGT’s targets regulate gene expression [17,18], metabolism [16,19,20], and signaling [21,22], consistent with OGT’s role in development and disease [23,24].

OGT stably interacts with and modifies all three TET proteins and its genome-wide distribution overlaps significantly with TETs [25-27]. Two studies in mouse embryonic stem cells (mESCs) have suggested that TET1 and OGT may be intimately linked in regulation of gene expression, as depleting either enzyme reduced the chromatin association of the other and affected expression of its target genes [25,28]. However, it is unclear to what extent these genome-wide changes are direct effects of perturbing the TET1-OGT interaction. Further work is necessary to uncover the biological importance of the partnership between TET1 and OGT.

In this work, we map the interaction between TET1 and OGT to a small C-terminal region of TET1, that is both necessary and sufficient to bind OGT. We show for the first time that OGT modifies the catalytic domain of TET1 *in vitro* and enhances its catalytic activity. Finally, we show that OGT levels correlate with TET activity in mESCs and use mutant TET1 to show that the TET1-OGT interaction promotes Tet1 function in the developing zebrafish embryo. Together these results suggest that OGT regulates TET1 activity, indicating that the TET1-OGT interaction may be two-fold in function – allowing TET1 to recruit OGT to specific genomic loci and allowing OGT to modulate TET1 activity.

## Materials and Methods

### Cell Culture

The mESC line LF2, and its derivatives, and a line expressing transgenic BirA-tag OGT [25] were routinely passaged by standard methods in KO-DMEM, 10% FBS, 2 mM glutamine, 1X non-essential amino acids, 0.1 mM b-mercaptoethanol and recombinant leukemia inhibitory factor. HEK293T were cultured in DMEM, 10% FBS, and 2 mM glutamine.

### Recombinant protein purification

Full-length human OGT in the pBJG vector was transformed into BL-21 DE3 E. coli. A liquid culture was grown in LB + 50ug/mL kanamycin at 37C until OD_600_ reached 1.0. IPTG was added to 1mM final and the culture was induced at 16C overnight. Cells were pelleted by centrifugation and resuspended in 5mL BugBuster (Novagen) + protease inhibitors (Sigma Aldrich) per gram of cell pellet. Cells were lysed on an orbital shaker for 20 minutes at room temperature. The lysate was clarified by centrifugation at 30,000g for 30 minutes at 4C. Clarified lysate was bound to Ni-NTA resin (Qiagen) at 4C and then poured over a disposable column. The column was washed with 6 column volumes of wash buffer 1 (20mM Tris pH 8, 1mM CHAPS, 10% glycerol, 5mM BME, 10mM imidazole, 250mM NaCl) followed by 6 column volumes of wash buffer 2 (wash buffer 1 with 50mM imidazole). The protein was eluted in 4 column volumes of elution buffer (20mM Tris pH 8, 1mM CHAPS, 5mM BME, 250mM imidazole, 250mM NaCl). Positive fractions were pooled and dialyzed into storage buffer (20mM Tris pH 8, 1mM CHAPS, 0.5mM THP, 10% glycerol, 150mM NaCl, 1mM EDTA), flash frozen in liquid nitrogen and stored at -80C in small aliquots.

Mouse TET1 catalytic domain (aa1367-2039) was expressed in sf9 insect cells according to the Bac-to-Bac Baculovirus Expression System. Constructs were cloned into the pFastBac HTA vector and transformed in DH10Bac E. coli for recombination into a bacmid. Bacmid containing the insert was isolated and used to transfect adherent sf9 cells for 6 days at 25C. Cell media (P1 virus) was isolated and used to infect 20mL of sf9 cells in suspension for 3 days. Cell media (P2 virus) was isolated and used to infect a larger sf9 suspension culture for 3 days. Cells were pelleted by centrifugation, resuspended in lysis buffer (20mM Tris pH 8, 1% Triton, 10% glycerol, 20mM imidazole, 50mM NaCl, 1mM MgCl_2_, 0.5mM TCEP, protease inhibitors, 2.5U/mL benzonase), and lysed by douncing and agitation at 4C for 1 hour. The lysate was clarified by centrifugation at 48,000g for 30 minutes at 4C and bound to Ni-NTA resin (Qiagen) at 4C, then poured over a disposable column. The column was washed with 5 column volumes of wash buffer (20mM Tris pH 8, 0.3% Triton, 10% glycerol, 20mM imidazole, 250mM NaCl, 0.5mM TCEP, protease inhibitors). The protein was eluted in 5 column volumes of elution buffer (20mM Tris pH 8, 250mM imidazole, 250mM NaCl, 0.5mM TCEP, protease inhibitors). Positive fractions were pooled and dialyzed overnight into storage buffer (20mM Tris pH 8, 150mM NaCl, 0.5mM TCEP). Dialyzed protein was purified by size exclusion chromatography on a 120mL Superdex 200 column (GE Healthcare). Positive fractions were pooled, concentrated, flash frozen in liquid nitrogen and stored at -80C in small aliquots.

### Overexpression in HEK293T cells and immunoprecipitation

Mouse Tet1 catalytic domain (aa1367-2039) and truncations and mutations thereof were cloned into the pcDNA3b vector. GFP fusion constructs were cloned into the pcDNA3.1 vector. Human OGT constructs were cloned into the pcDNA4 vector. Plasmids were transiently transfected into adherent HEK293T cells at 70-90% confluency using the Lipofectamine 2000 transfection reagent (ThermoFisher) for 1-3 days.

Full-length mouse Tet1 and mutations thereof were cloned into the pCAG vector. Plasmids were transiently transfected into adherent HEK293T cells at 70-90% confluency using the PolyJet transfection reagent (SignaGen) for 1-3 days.

Transiently transfected HEK293T cells were harvested, pelleted, and lysed in IP lysis buffer (50mM Tris pH 8, 200mM NaCl, 1% NP40, 1x HALT protease/phosphatase inhibitors). For pulldown of FLAG-tagged constructs, cell lysate was bound to anti-FLAG M2 magnetic beads (Sigma Aldrich) at 4C. For pulldown of GFP constructs, cell lysate was bound to magnetic protein G dynabeads (ThermoFisher) conjugated to the JL8 GFP monoclonal antibody (Clontech) at 4C. Beads were washed 3 times with IP wash buffer (50mM Tris pH 8, 200mM NaCl, 0.2% NP40, 1x HALT protease/phosphatase inhibitors). Bound proteins were eluted by boiling in SDS sample buffer.

### *In vitro* transcription/translation and immunoprecipitation

GFP fused to TET C-terminus peptides were cloned into the pcDNA3.1 vector and transcribed and translated *in vitro* using the TNT Quick Coupled Transcription/Translation System (Promega).

For immunoprecipitation, recombinant His-tagged OGT was coupled to His-Tag isolation dynabeads (ThermoFisher). Beads were bound to *in vitro* translation extract diluted 1:1 in binding buffer (40mM Tris pH 8, 200mM NaCl, 40mM imidazole, 0.1% NP40) at 4C. Beads were washed 3 times with wash buffer (20mM Tris pH 8, 150mM NaCl, 20mM imidazole, 0.1% NP40). Bound proteins were eluted by boiling in SDS sample buffer.

### Recombinant protein binding assay

20uL reactions containing 2.5uM rOGT and 2.5uM rTET1 CD wt or D2018A were assembled in binding buffer (50mM Tris pH 7.5, 100mM NaCl, 0.02% Tween-20) and pre-incubated at room temperature for 15 minutes. TET1 antibody (Millipore 09-872) was bound to magnetic Protein G Dynabeads (Invitrogen), and beads added to reactions following pre-incubation. Reactions were bound to beads for 10 minutes at room temperature. Beads were washed 3 times with 100uL binding buffer, and bound proteins were recovered by boiling in SDS sample buffer and analyzed by SDS-PAGE and coomassie stain.

### Western blots

For western blot, proteins were separated on a denaturing SDS-PAGE gel and transferred to PVDF membrane. Membranes were blocked in PBST + 5% nonfat dry milk at room temp for >10 minutes or at 4C overnight. Primary antibodies used for western blot were: FLAG M2 monoclonal antibody (Sigma Aldrich F1804), OGT polyclonal antibody (Santa Cruz sc32921), OGT monoclonal antibody (Cell Signaling D1D8Q), His6 monoclonal antibody (Thermo MA1-21315), and JL8 GFP monoclonal antibody (Clontech). Secondary antibodies used were goat anti-mouse HRP and goat anti-rabbit HRP from BioRad. Blots were incubated with Pico Chemiluminescent Substrate (ThermoFisher) and exposed to film in a dark room.

### Slot blot

Prior to dilution of genomic DNA samples, biotinylated *E. coli* gDNA was added as a loading control (see below). DNA samples were denatured in 400mM NaOH + 10mM EDTA by heating to 95C for 10 minutes. Samples were placed on ice and neutralized by addition of 1 volume of cold NH_4_OAc pH 7.2. DNA was loaded onto a Hybond N+ nylon membrane (GE) by vacuum using a slot blot apparatus. The membrane was dried at 37C and DNA was covalently linked to the membrane by UV crosslinking (700uJ/cm^2^ for 3 minutes). Antibody binding and signal detection were performed as outlined for western blotting. Primary antibodies used were 5mC monoclonal antibody (Active Motif 39649) and 5hmC monoclonal antibody (Active Motif 39791).

For the loading control, membranes were analyzed using the Biotin Chromogenic Detection Kit (Thermo Scientific) according to the protocol. Briefly, membranes were blocked, probed with streptavidin conjugated to alkaline phosphatase (AP), and incubated in the AP substrate BCIP-T (5-bromo-4-chloro-3-indolyl phosphate, p-toluidine salt). Cleavage of BCIP-T causes formation of a blue precipitate.

### Preparation of lambda DNA substrate

Linear genomic DNA from phage lambda (dam-, dcm-) containing 12bp 5’ overhangs was purchased from Thermo Scientific. Biotinylation was performed by annealing and ligating a complementary biotinylated DNA oligo. Reactions containing 175ng/uL lambda DNA, 2uM biotinylated oligo, and 10mM ATP were assembled in 1x T4 DNA ligase buffer, heated to 65C, and cooled slowly to room temperature to anneal. 10uL T4 DNA ligase was added and ligation was performed overnight at room temperature. Biotinylated lambda DNA was purified by PEG precipitation. To a 500uL ligation reaction, 250uL of PEG8000 + 10mM MgCl_2_ was added and reaction was incubated at 4C overnight with rotation. The next day DNA was pelleted by centrifugation at 14,000g at 4C for 5 minutes. Pellet was washed with 1mL of 75% ethanol and resuspended in TE.

Biotinylated lambda DNA was methylated using M.SssI CpG methyltransferase from NEB. 20uL reactions containing 500ng lambda DNA, 640uM S-adenosylmethionine, and 4 units methyltransferase were assembled in 1x NEBuffer 2 supplemented with 20mM Tris pH 8 and incubated at 37C for 1 hour. Complete methylation was confirmed by digestion with the methylation-sensitive restriction enzyme BstUI from NEB.

### *In vitro* TET1 CD O-GlcNAcylation

*In vitro* modification of rTET1 CD with rOGT was performed as follows: 10uL reactions containing 1uM rTET1 CD, 10uM rOGT, and 2.5mM UDP-GlcNAc were assembled in reaction buffer (50mM Tris pH 7.5, 12.5mM MgCl_2_, 2% glycerol, 1mM DTT) and incubated at 37C for 30-60 minutes.

### *In vitro* TET1 CD activity assays

20uL reactions containing 100ng biotinylated, methylated lambda DNA, rTET1 CD (from frozen aliquots or from *in vitro O*-GlcNAcylation reactions), and TET cofactors (1mM alpha-ketoglutarate, 2mM ascorbic acid, 100uM ferrous ammonium sulfate) were assembled in reaction buffer (50mM HEPES pH 6.5, 100mM NaCl) and incubated at 37C for 1 hour. Reactions were stopped by incubation at 65C for 5 minutes, then cooled slowly to room temperature to allow re-annealing of DNA. DNA was digested to fragments <10kb and purified using DNA clean and concentrator kit (Zymo). 5hmC content was analyzed by slot blot.

### Generation of mouse embryonic stem cell lines

mESC lines (Fig. S4A, B) were derived using CRISPR-Cas9 genome editing. A guide RNA to the Tet1 3’UTR was cloned into the px459-Cas9-2A-Puro plasmid using published protocols [29] with minor modifications. Templates for homology directed repair were amplified from Gene Blocks (IDT) (Tables S1 and S2). Plasmid and template were co-transfected into LF2 mESCs using FuGENE HD (Promega) according to manufacturer protocol. After two days cells were selected with puromycin for 48 hours, then allowed to grow in antibiotic-free media. Cells were monitored for green or red fluorescence (indicating homology directed repair) and fluorescent cells were isolated by FACS 1-2 weeks after transfection. All cell lines were propagated from single cells and correct insertion was confirmed by PCR genotyping (Fig. S4A, B, Table S1).

### Immunofluorescence

For IF experiments, cells were cytospun onto glass slides at 800rpm for 3 minutes. For 5mC or 5hmC staining alone, cells were fixed in 3:1 methanol:acetic acid for 5 minutes. For co-staining for 5hmC and FLAG, cells were fixed in 4% paraformaldehyde for 10 minutes. Fixed slides were washed with PBS + 0.01% Tween-20 (PBST) and stored in 70% ethanol at -20C until use.

Fixed slides were incubated in 1M HCl at 37C for 45 minutes to denature chromatin, then neutralized in 100mM Tris pH 7.6 at room temperature for 10 minutes. Slides were washed twice in PBST for 5 minutes, then blocked in IF blocking buffer (PBS + 5% goat serum,0.2% fish skin gelatin, 0.2% Tween-20) at room temperature for 1 hour. Primary antibodies were diluted in blocking buffer and incubated on slides at 4C overnight. Primary antibodies used were FLAG M2 monoclonal antibody (Sigma Aldrich F1804), 5mC monoclonal antibody (Active Motif 39649), and 5hmC polyclonal antibody (Active Motif 39791). Slides were washed twice in PBST for 5 minutes, then incubated with secondary antibodies diluted in IF blocking buffer. Secondary antibodies used were Alexa488-conjugated goat anti-rabbit IgG (Jackson 711-545-152), Cy3-conjugated goat anti-rabbit IgG (Jackson 715-165-152), and Cy3-conjugated goat anti-mouse IgG (Jackson 715-165-150). Slides were washed three times in PBST for 5 minutes with DAPI added to the second wash (final concentration 100ng/mL). Slides were then mounted using prolong gold antifade (Molecular Probes P36930) and imaged.

### mRNA rescue experiments

Zebrafish husbandry was conducted under full animal use and care guidelines with approval by the Memorial Sloan-Kettering animal care and use committee. For mRNA rescue experiments, mTET1D2018A and mTET1wt plasmids were linearized by NotI digestion. Capped RNA was synthesized using mMessage mMachine (Ambion) with T7 RNA polymerase. RNA was injected into one-cell-stage embryos derived from tet2^mk17/mk17^, tet3^mk18/+^ intercrosses at the concentration of 100pg/embryo [30]. Injected embryos were raised under standard conditions at 28.5°C until 30 hours post-fertilization (hpf) at which point they were fixed for *in situ* hybridization using an antisense probe for *runx1*. The *runx1* probe is described in [31]; *in situ* hybridization was performed using standard methods [32]. *tet2/3* double mutants were identified based on morphological criteria and mutants were confirmed by PCR genotyping after in situ hybridization using previously described primers [30].

For sample size estimation for rescue experiments, we assume a background mean of 20% positive animals in control groups. We anticipate a significant change would result in at least a 30% difference between the experimental and control means with a standard deviation of no more than 10. Using the 1-Sample Z-test method, for a specified power of 95% the minimum sample size is 4. Typically, zebrafish crosses generate far more embryos than required. Experiments are conducted using all available embryos. The experiment is discarded if numbers for any sample are below this minimum threshold when embryos are genotyped at the end of the experimental period. Results from five independent experiments were reported; p values were derived from the unpaired two-tailed *t* test.

For the dot blot, genomic DNA was isolated from larvae at 30hpf by phenol-chloroform extraction and ethanol precipitation. Following RNase treatment and denaturation, 2-fold serially diluted DNA was spotted onto nitrocellulose membranes. Cross-linked membranes were incubated with 0.02% methylene blue to validate uniform DNA loading. Membranes were blocked with 5% BSA and incubated with anti-5hmC antibody (1:10,000; Active Motif) followed by a horseradish peroxidase-conjugated antibody (1:15,000; Active Motif). Signal was detected using the ECL Prime Detection Kit (GE).

### Reproducibility and Rigor

All immunostaining, IP-Westerns, and genomic DNA slot blot data are representative of at least three independent biological replicates (experiments carried out on different days with a different batch of HEK293T cells or mESCs). For targeted mESC lines, three independently derived lines for each genotype were assayed in at least two biological replicates. For *in vitro* activity and binding assays using recombinant proteins (representing multiple protein preparations), data represent at least three technical replicates (carried out on multiple days). The zebrafish rescue experiment was performed five times (biological replicates), with dot blots carried out three times. We define an outlier as a result in which all the controls gave the expected outcome but the experimental sample yielded an outcome different from other biological or technical replicates. There were no outliers or exclusions.

## Results

### OGT enhances TET activity *in vitro* and in cells

TETs have been implicated in localizing OGT to chromatin, however a role for OGT in regulation of TETs has not been fully explored. To examine whether OGT can regulate activity of a TET protein, we employed recombinant enzymes to assess the effects of OGT on TET1 activity. We purified recombinant mouse TET1 catalytic domain (rTET1 CD, aa1367-2039) from sf9 cells and recombinant human OGT (rOGT; identical to mouse at 1042 of 1046 residues) from *E. coli* (Fig. S1). rTET1 CD catalyzed formation of 5hmC on an *in vitro* methylated lambda DNA substrate, dependent on the presence of the TET cofactors alpha-ketoglutarate, Fe^2+^, and ascorbic acid (Fig. S2). Incubation of rTET1 CD with rOGT and OGT’s cofactor UDP-GlcNAc resulted in *O*-GlcNAcylation of rTET1 CD (Fig. 1A). Modified rTET1 CD produced more 5hmC than unmodified rTET1 CD (Fig. 1B, C), indicating that OGT stimulates the activity of the TET1 CD *in vitro*.

**Fig. 1:**
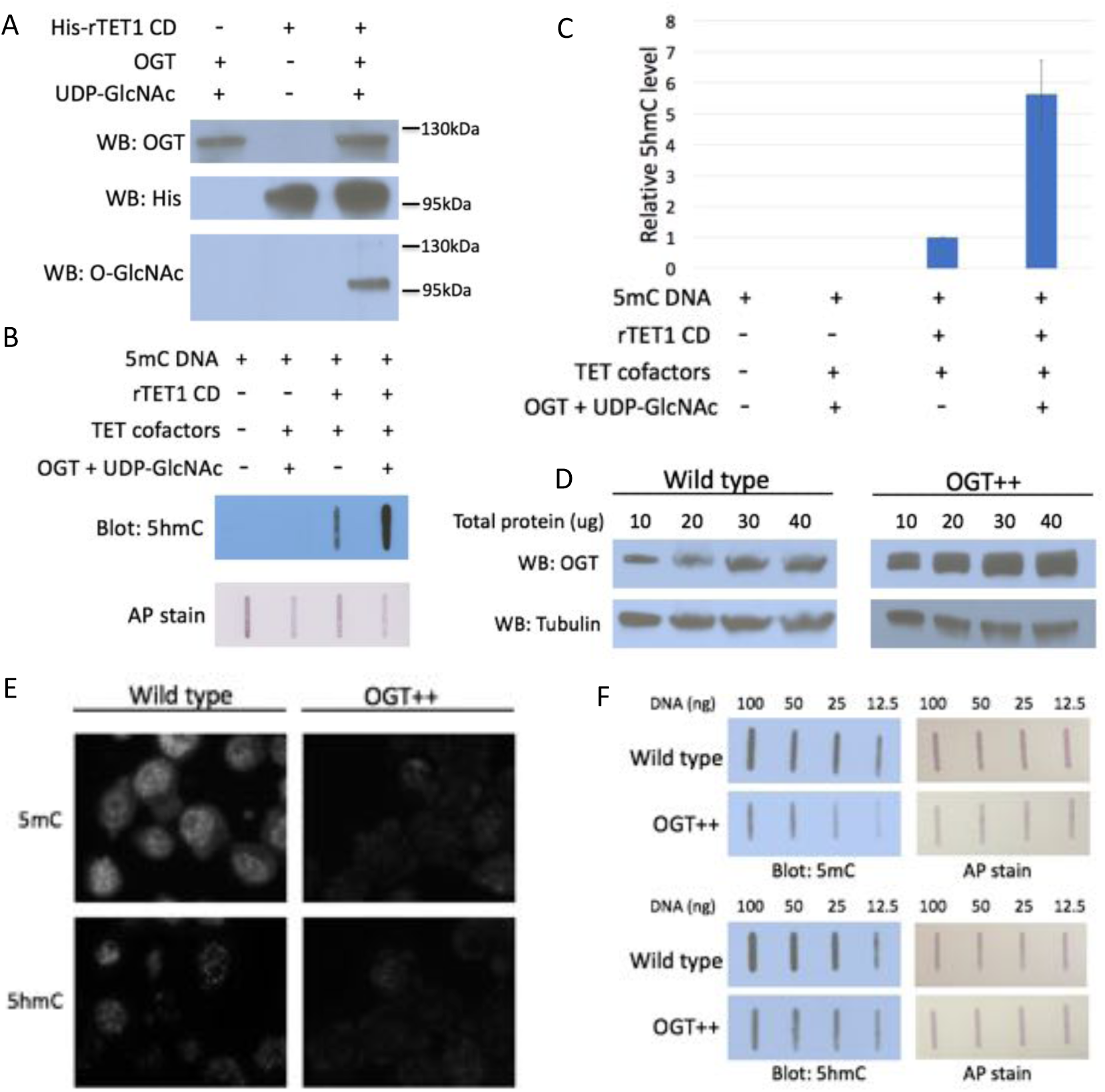
OGT enhances TET activity *in vitro* and in cells. A) Western blot for OGT, rTET1 CD, and *O*-GlcNAc in *in vitro O*-GlcNAcylation reaction demonstrating UDP-GlcNAc dependent TET1 CD *O*-GlcNAcylation. B) 5hmC slot blot of biotinylated 5mC containing lambda DNA from rTET1 CD activity assays. Alkaline phosphatase staining was used to detect biotin as a loading control. C) Quantification of 5hmC levels from four slot blots. All results normalized to TET1 CD without UDP-GlcNAc and OGT. D) Western blot of 10-40ug lysate from wild type and OGT overexpressing ( OGT++) mESCs, showing increased OGT levels in OGT++ mESCs. E) Immunofluorescence comparing 5mC (upper) and 5hmC (lower) levels in wild type vs. OGT++ mESCs. F) Slot blots for 5mC and 5hmC of 12.5-100 ng genomic DNA from wild type and OGT++ mESCs. Equal amounts of biotinylated plasmid DNA were added to each gDNA dilution and alkaline phosphatase staining was used to detect biotin as a loading control.

To examine whether OGT affects TET activity in a more biological context, we assessed the effects of altering OGT levels in mESCs. These cells express TET1 and TET2 [33] and the proteins that remove 5fC and 5caC and replace them with unmodified cytosine. Thus increased TET activity should manifest as decreased 5mC and 5hmC. We examined levels of 5mC and 5hmC in mESCs overexpressing OGT (Fig. 1D) by immunofluorescence (Fig. 1E) and by slot blot of genomic DNA (Fig. 1F). These experiments clearly show that overexpression of OGT reduces the levels of both 5mC and 5hmC, consistent with increased TET activity oxidizing 5hmC to 5fC and 5caC. These results are consistent with OGT stimulation of TET activity in mESCs.

### A short C-terminal region of TET1 is necessary for binding to OGT

To examine whether OGT regulates TET1 activity *in vivo*, it is necessary to specifically disrupt the TET1-OGT interaction. All three TETs interact with OGT via their catalytic domains [26,27,34]. We sought to identify the domain within the TET1 CD responsible for binding to OGT. The TET1 CD consists of a cysteine rich N-terminal region necessary for co-factor and substrate binding, a catalytic fold consisting of two lobes separated by a spacer of unknown function, and a short C-terminal region also of unknown function (Fig. 2A). We transiently transfected HEK293T cells with FLAG-tagged mouse TET1 CD constructs bearing deletions of each of these regions, some of which failed to express (Fig. 2B). Because HEK293T cells have low levels of endogenous OGT, we also co-expressed myc-tagged human OGT. TET1 constructs were immunoprecipitated (IPed) using FLAG antibody and analyzed for interaction with OGT. We found that deletion of only the 45 residue C-terminus of TET1 (hereafter TET1 C45) prevented detectable interaction with OGT (Fig. 2C, TET1 CD del. 4). To exclude the possibility that this result is an artifact of OGT overexpression, we repeated the experiment overexpressing only TET1. TET1 CD, but not TET1 CD ΔC45, interacted with endogenous OGT, confirming that the C45 is necessary for this interaction (Fig. S3).

**Fig. 2:**
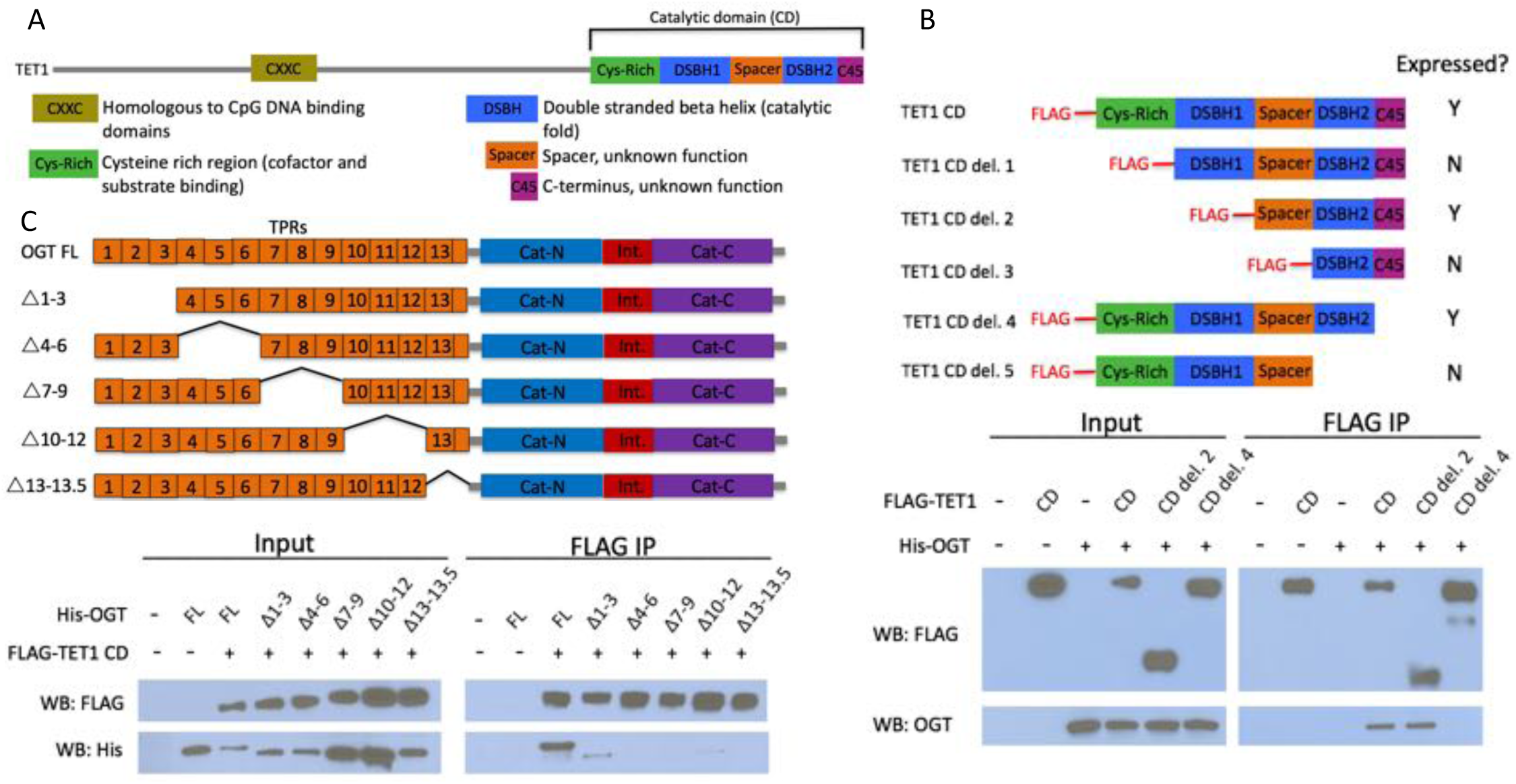
The short TET1 C-terminus is required for interaction with OGT. A) Domain architecture of TET1. B) Diagram of FLAG-tagged TET1 CD constructs expressed in HEK293T cells (upper). FLAG and OGT western blot of inputs and FLAG IPs from HEK293T cells transiently expressing FLAG-TET1 CD truncations and His-OGT ( lower). C) Diagram of His-tagged OGT constructs expressed in HEK293T cells (upper). FLAG and His western blot of input and FLAG IPs from HEK293T cells transiently expressing FLAG-TET1 CD and His-OGT TPR deletions (lower).

OGT has two major domains: the N-terminus consists of 13.5 tetratricopeptide repeat (TPR) protein-protein interaction domains, and the C-terminus contains the bilobed catalytic domain (Fig. 2D). We made internal deletions of several sets of TPRs to ask which are responsible for binding to the TET1 CD. We co-transfected HEK293T cells with FLAG-TET1 CD and His6-tagged OGT constructs and performed FLAG IP and western blot as above. We found that all the TPR deletions tested impaired the interaction with TET1 CD, with deletion of TPRs 7-9, 10-12, or 13-13.5 being most severe (Fig. 2E). This result suggests that all of OGT’s TPRs may be involved in binding to the TET1 CD, or that deletion of a set of TPRs disrupts the overall structure of the repeats in a way that disfavors binding.

### Conserved residues in the TET1 C45 are necessary for the TET1-OGT interaction

An alignment of the TET1 C45 region with the C-termini of TET2 and TET3 revealed several conserved residues (Fig. 3A). We mutated clusters of three conserved residues in the TET1 C45 of FLAG-tagged TET1 CD (Fig. 3B) and co-expressed these constructs with myc-OGT in HEK293T cells. FLAG pulldowns revealed that two sets of point mutations disrupted the interaction with OGT: mutation of D2018, V2021, and T2022, or mutation of V2021, T2022, and S2024 (Fig. 3C, mt1 and mt2). These results suggested that the residues between D2018 and S2024 are crucial for the interaction between TET1 and OGT. We mutated these residues individually and found that D2018A eliminated detectable interaction between FLAG-tagged TET1 CD and myc-OGT (Fig. 3D).

**Fig. 3:**
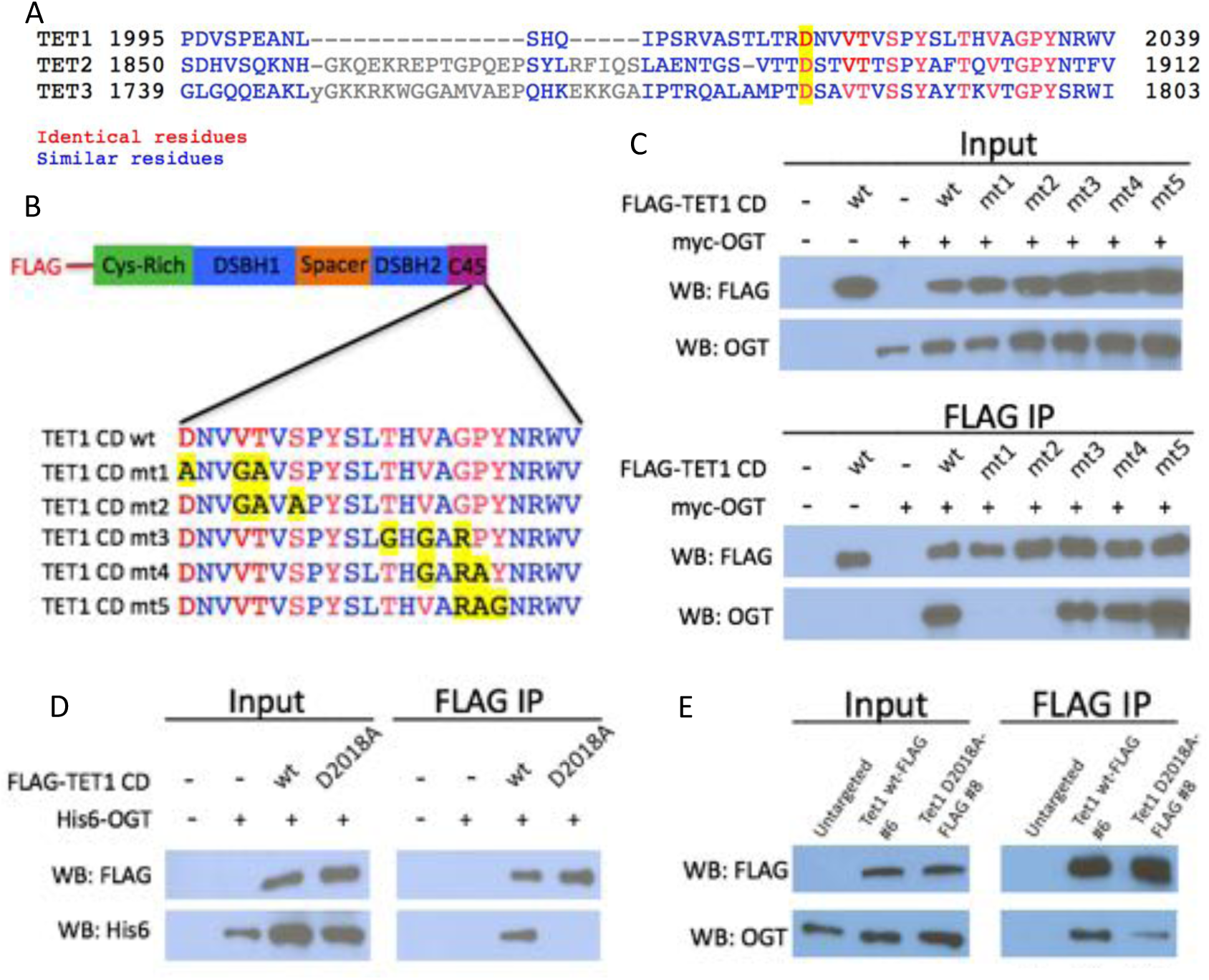
Conserved residues in the TET1 C45 are necessary for the TET1-OGT interaction. A) Alignment of the C-termini of mouse TETs 1, 2, and 3. A conserved aspartate residue mutated in D and E is highlighted. B) Diagram of FLAG-tagged TET1 CD constructs expressed in HEK293T cells. C) FLAG and OGT western blot of inputs and FLAG IPs from HEK293T cells transiently expressing FLAG-TET1 CD triple point mutants and His-OGT. D) FLAG and OGT western blot of inputs and FLAG IPs from HEK293T cells transiently expressing His-OGT and F LAG-TET1 CD or F LAG-TET1 CD D2018A. E) FLAG and OGT western blot of inputs and FLAG IPs from mESCs in which a FLAG tag or a FLAG tag and the D2018A mutation were introduced into both copies of *Tet1* (Fig. S4).

To examine whether D2018 was necessary for the TET1-OGT interaction in the context of full length TET1, we generated a D2018A mutation in both copies of the *Tet1* gene in mESCs (Fig. S4A, B). A FLAG tag was also introduced onto the C-terminus of wild type or D2018A mutant TET1. FLAG pulldowns revealed that the D2018A mutation reduced, but did not eliminate, co-IP of OGT with TET1 (Fig. 3E). This reduction of OGT was consistent in multiple experiments and across multiple independently derived cell lines (Fig. S4C). In mESCs, TET1 and OGT are found together in high molecular weight complexes [25] and interactions of TET1 and OGT with other proteins in these complexes may be sufficient to support some TET1-OGT interaction when D2018 is mutated (see discussion).

### The TET1 C-terminus is sufficient for binding to OGT

Having shown that the TET1 C45 is necessary for the interaction with OGT, we next examined if it is also sufficient to bind OGT. We fused the TET1 C45 to the C-terminus of GFP (Fig. 4A) and investigated its interaction with OGT. We transiently transfected GFP or GFP-TET1 C45 into HEK293T cells and pulled down with a GFP antibody. We found that GFP-TET1 C45, but not GFP alone, bound OGT (Fig. 4B), indicating that the TET1 C45 is sufficient for interaction with OGT.

**Fig. 4:**
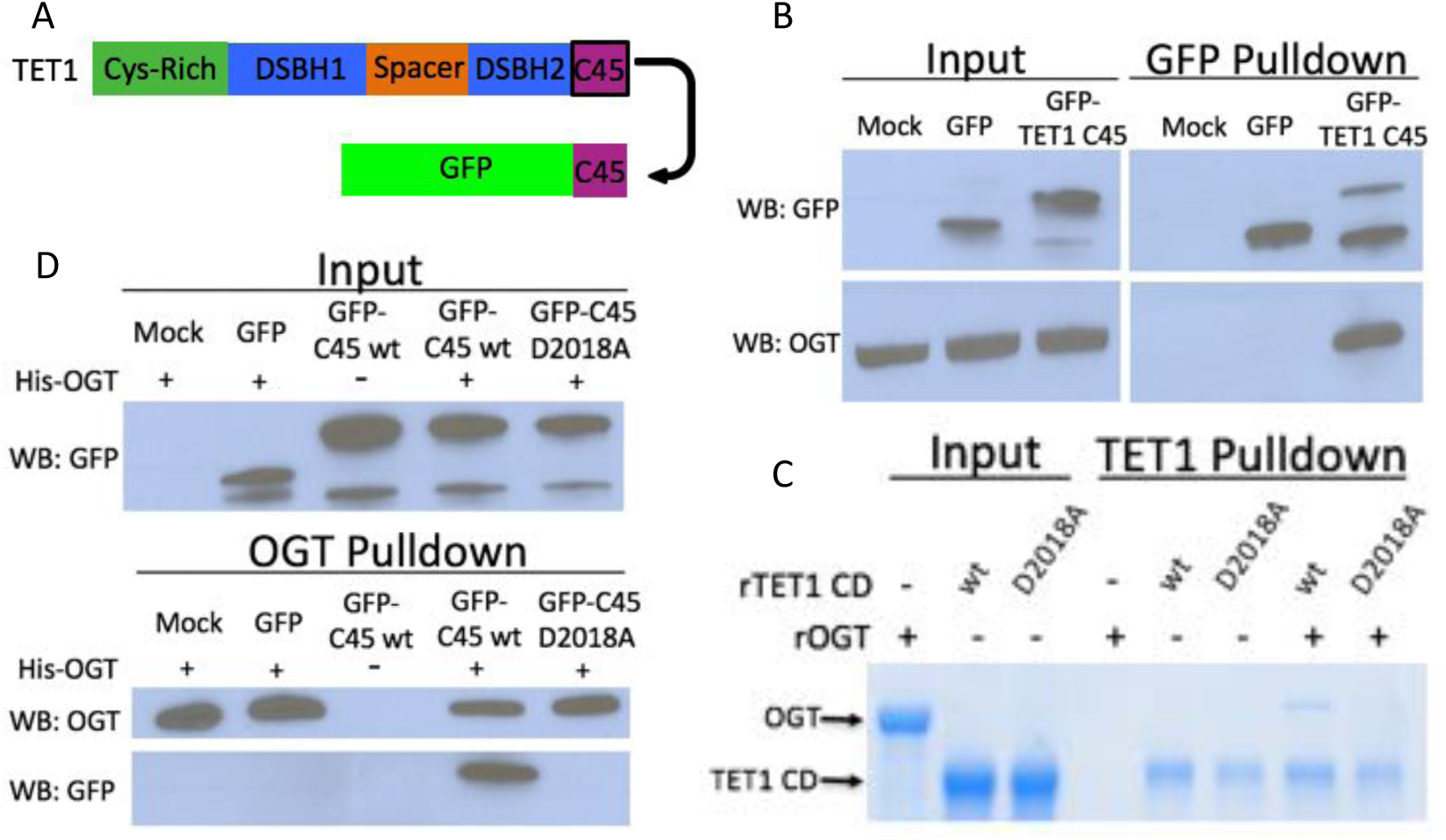
The TET1 C45 is sufficient for interaction with OGT in cells and in vitro. A) Schematic of the TET1 C45 fusion to the C-terminus of GFP. B) GFP and OGT western blot of inputs and GFP IPs from HEK293T cells transiently expressing GFP or GFP-TET1 C45. C) Coomassie stained protein gel of inputs and TET1 IPs from *in vitro* binding reactions containing rOGT and rTET1 CD wild type or D2018A. D) GFP and OGT western blot of inputs and OGT IPs from *in vitro* binding reactions containing rOGT and *in vitro* translated GFP constructs.

To determine if the interaction between TET1 CD and OGT is direct, we employed recombinant proteins in pulldown assays using beads conjugated to a TET1 antibody. We found that wild-type rTET1 CD, but not beads alone, pulled down OGT, indicating a direct interaction between these proteins (Fig. 4C). rTET1 CD D2018A did not pull down rOGT, consistent with our mutational analysis in cells. Then we used an *in vitro* transcription/translation extract to produce GFP and GFP-TET1 C45, incubated each with rOGT, and found that the TET1 C45 is sufficient to confer binding to rOGT (Fig. 4D). The D2018A mutation was also sufficient in this context to prevent OGT binding (Fig. 4D), consistent with the behavior of this mutant in cells. Together these results indicate that the TET1-OGT interaction is direct and mediated by the TET1 C45.

### TET1 D2018A is active in HEK293T cells

The D2018A mutation lies outside the catalytic fold, suggesting that this mutation is unlikely to affect TET1 activity. Consistent with this suggestion, in human TET2 the region homologous to TET1 C45 is dispensable for catalytic activity *in vitro* [35]. To test whether full length TET1 D2018A is catalytically active, we examined its activity in cells. FLAG-tagged full-length wild type or D2018A TET1 were transfected into HEK293T cells, which express low levels of OGT and of all three TETs, facilitating a more direct comparison of the two TET1 constructs. Western blot confirmed that the two proteins were expressed at comparable levels (Fig. 5A). Immunostaining for FLAG and 5hmC showed that wild type and D2018A TET1 both exhibited nuclear localization (Fig. 5B). While mock transfected HEK293T cells showed very low levels of 5hmC, consistent with their low expression of TETs and OGT, wild type or D2018A TET1 each caused increased production of 5hmC (Fig. 5B). A slot blot assay on genomic DNA from these cells confirmed that comparable levels of 5hmC were observed in wild type and D2018A TET1 transfections (Fig. 5C). These results indicate that the D2018A mutation does not notably impair TET1’s catalytic activity.

**Fig. 5:**
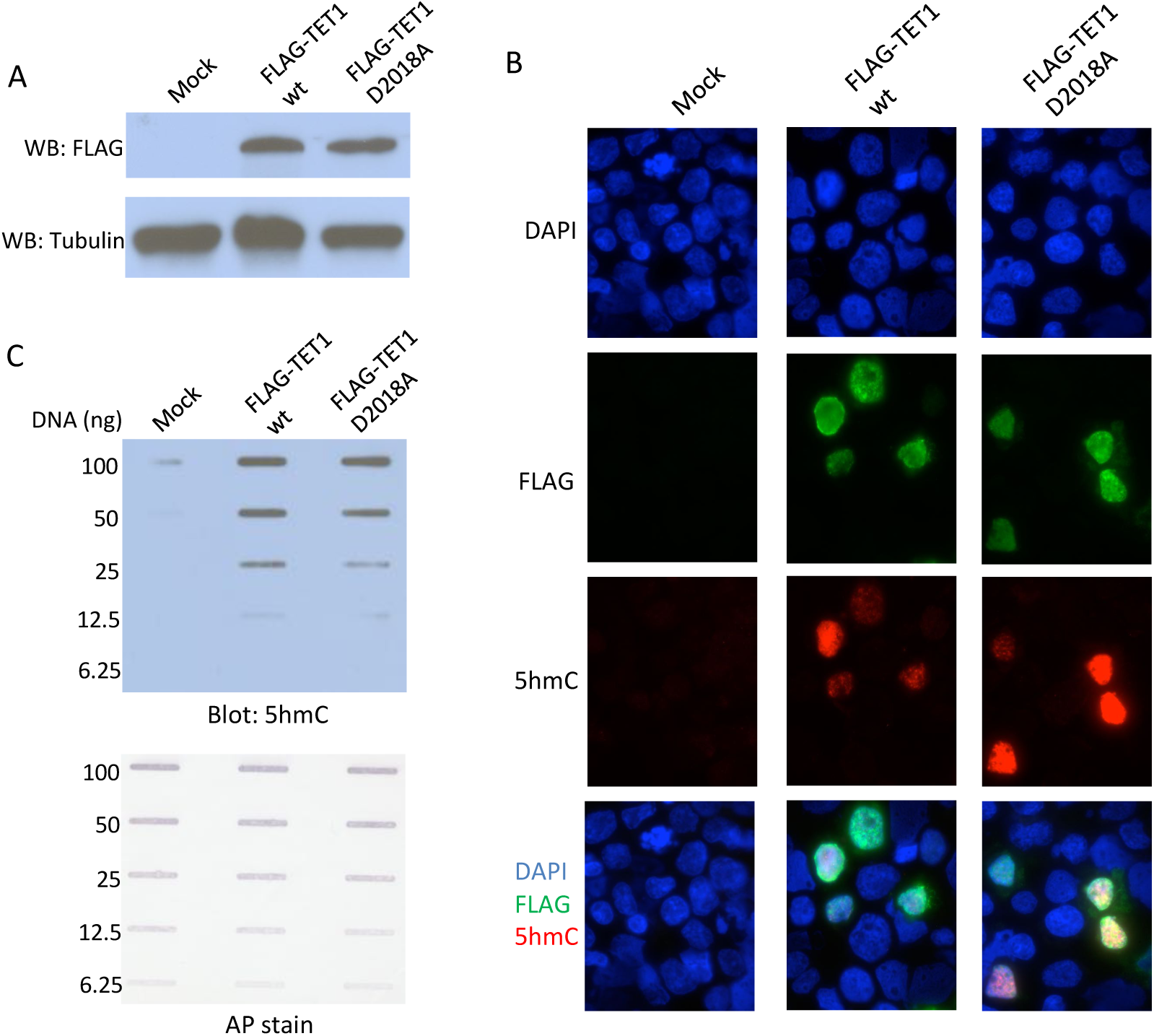
TET1 D2018A is active in HEK293T cells. A) FLAG western blot of HEK293T cell lysates transiently expressing FLAG-TET1 wild type or D2018A. Western blot for Tubulin was performed as a loading control. B) Immunofluorescence for FLAG (green) and 5hmC (red) in HEK293T cells transiently expressing FLAG-TET1 wild type or D2018A. DAPI ( blue) was used to stain nuclei. C) 5hmC slot blot of 6.25-100ng genomic DNA from HEK293T cells transiently expressing FLAG-TET1 wild type or D2018A. Equal amounts of biotinylated plasmid DNA were added to each gDNA stock and diluted across the dilution series. Alkaline phosphatase staining was used to detect biotin as a loading and dilution control.

### The TET-OGT interaction promotes Tet1 function in the zebrafish embryo

We used zebrafish as a model system to ask whether the D2018A mutation affects TET function during development. Deletion analysis of *tets* in zebrafish showed that Tet2 and Tet3 are the most important in development, while Tet1 contribution is relatively limited [30]. Deletion of both *tet2* and *tet3* (*tet2/3^DM^*) causes a severe decrease in 5hmC levels accompanied by larval stage lethality owing to a number of abnormalities including defects in hematopoietic stem cell (HSC) production. A critical transcription factor, Runx1, marks HSCs in the dorsal aorta of wild-type embryos, but is largely absent in this region of *tet2/3^DM^* embryos. 5hmC levels and *runx1* expression are rescued by injection of human TET2 or TET3 mRNA into one-cell-stage embryos [30].

Given strong sequence conservation among vertebrate TET/Tet proteins, we asked if over expression of Tet1 mRNA could also rescue HSCs in *tet2/3^DM^* zebrafish embryos and if this rescue is OGT interaction-dependent. To this end, *tet2/3^DM^* embryos were injected with wild type or D2018A mutant encoding Tet1 mRNA at the one cell stage. At 30 hours post fertilization (hpf) embryos were fixed and the presence of *runx1* positive HSCs in the dorsal aorta was assessed by *in situ* hybridization (Fig. 6A). Tet1 wild type mRNA significantly increased the percentage of embryos with high *runx1*, while Tet1 D2018A mRNA failed to rescue *runx1* positive cells (Fig. 6A-B). We also performed dot blots with genomic DNA from these embryos to measure levels of 5hmC (Fig. 6C). Embryos injected with wild type Tet1 mRNA showed a modest increase in 5hmC relative to uninjected *tet2/3^DM^* embryos, while injection of TET1 D2018A mRNA did not show an increase. These results suggest that the OGT-TET1 interaction promotes both TET1’s catalytic activity and its ability to rescue *runx1* expression in this system.

**Fig. 6:**
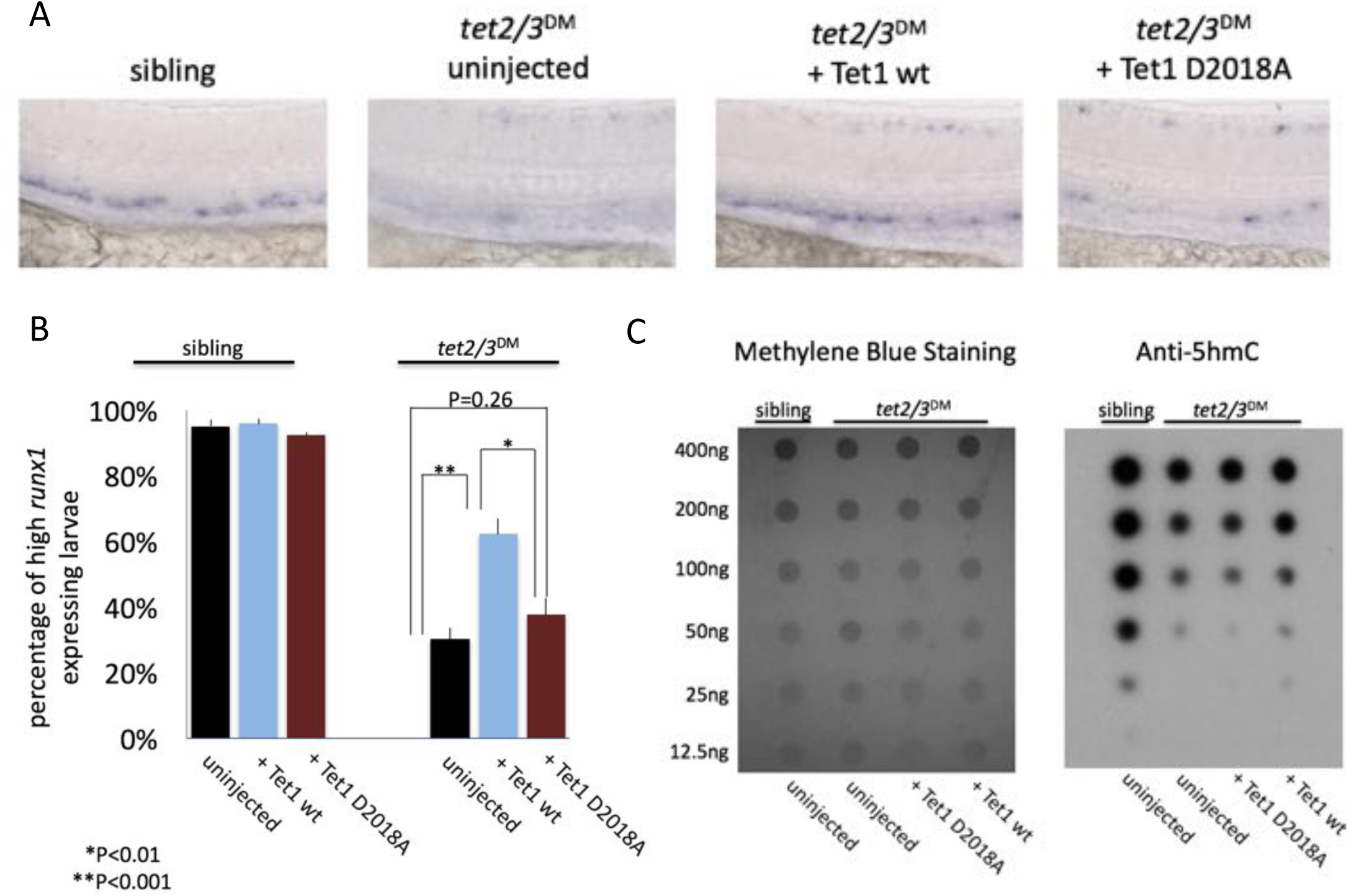
The interaction with OGT is required for TET1 activity in the zebrafish embryo. A) Representative images of *runx1* labeling in the dorsal aorta of wild type or *tet2/3^DM^* zebrafish embryos, uninjected or injected with mRNA encoding Tet1 wild type or D2018A, B) Percentage of embryos with high *runx1* expression along the dorsal aorta. C) 5hmC dot blot of genomic DNA from wild type or *tet2/3^DM^* zebrafish embryos injected with Tet1 wild type or D2018A mRNA. Methylene blue was used as a loading control.

## Discussion

### OGT stimulation of TET activity

Our results show for the first time that OGT stimulates the activity of a TET protein *in vitro*. The mechanism underlying this stimulation is unclear; it may be the result of the protein-protein interaction, the *O*-GlcNAc modification, or both. Further experiments uncoupling these elements will be informative.

Our data demonstrate that OGT stimulates TET1 activity *in vitro* and are consistent with a role for OGT in TET1 regulation *ex vivo* and *in vivo*. OGT also directly interacts with TET2 and TET3, suggesting that it may regulate all three TET proteins. Notably, although all three TETs possess highly similar catalytic folds and the same catalytic activity *in vitro*, they show a number of differences that are likely to determine their biological role. Different TET proteins are expressed in different cell types and at different stages of development [36-39]. TET1 and TET2 appear to target different genomic regions [40] and to promote different pluripotent states in mESCs [41]. The mechanisms responsible for these differences are not well understood. We suggest that OGT is a strong candidate for regulation of TET enzymes.

### A unique OGT interaction domain?

We identified a 45-amino acid domain of TET1 that is both necessary and sufficient for binding of OGT. To our knowledge, this is the first time that a small protein domain has been identified that confers stable binding to OGT. The vast majority of OGT targets do not bind to OGT tightly enough to be detected in co-IP experiments, suggesting that OGT’s interaction with TET proteins is unusually strong. For determination of the crystal structure of the human TET2 catalytic domain in complex with DNA, the corresponding C-terminal region had to be deleted [35], suggesting that it may be unstructured. When bound to OGT this domain may become structured, and structural studies of OGT bound to C45 could shed light on what features make this domain uniquely able to interact stably with OGT and how OGT may stimulate TET1 activity.

An alternative or additional role for the stable TET-OGT interaction may be recruitment of OGT to chromatin by TET proteins. Loss of TET1 causes loss of OGT from chromatin [25] and induces similar changes in transcription in both wild-type mESCs and mESCs lacking DNA methylation [42]. This raises the possibility that TET proteins may recruit OGT to chromatin to regulate gene expression independent of 5mC oxidation. Consistent with this possibility, OGT modifies many transcription factors and chromatin regulators in mESCs [43](Fig. 7). Thus it may be that the stable OGT-TET1 interaction promotes both regulation of TET1 activity by *O*-GlcNAcylation as well as recruitment of OGT to chromatin. Notably, our results show that TET1 D2018A does not rescue 5hmC levels in *tet2/3^DM^* zebrafish embryos to the same extent as the wild type protein, suggesting that at least part of the role of the OGT-TET1 interaction *in vivo* is regulation of TET1 activity.

**Fig. 7:**
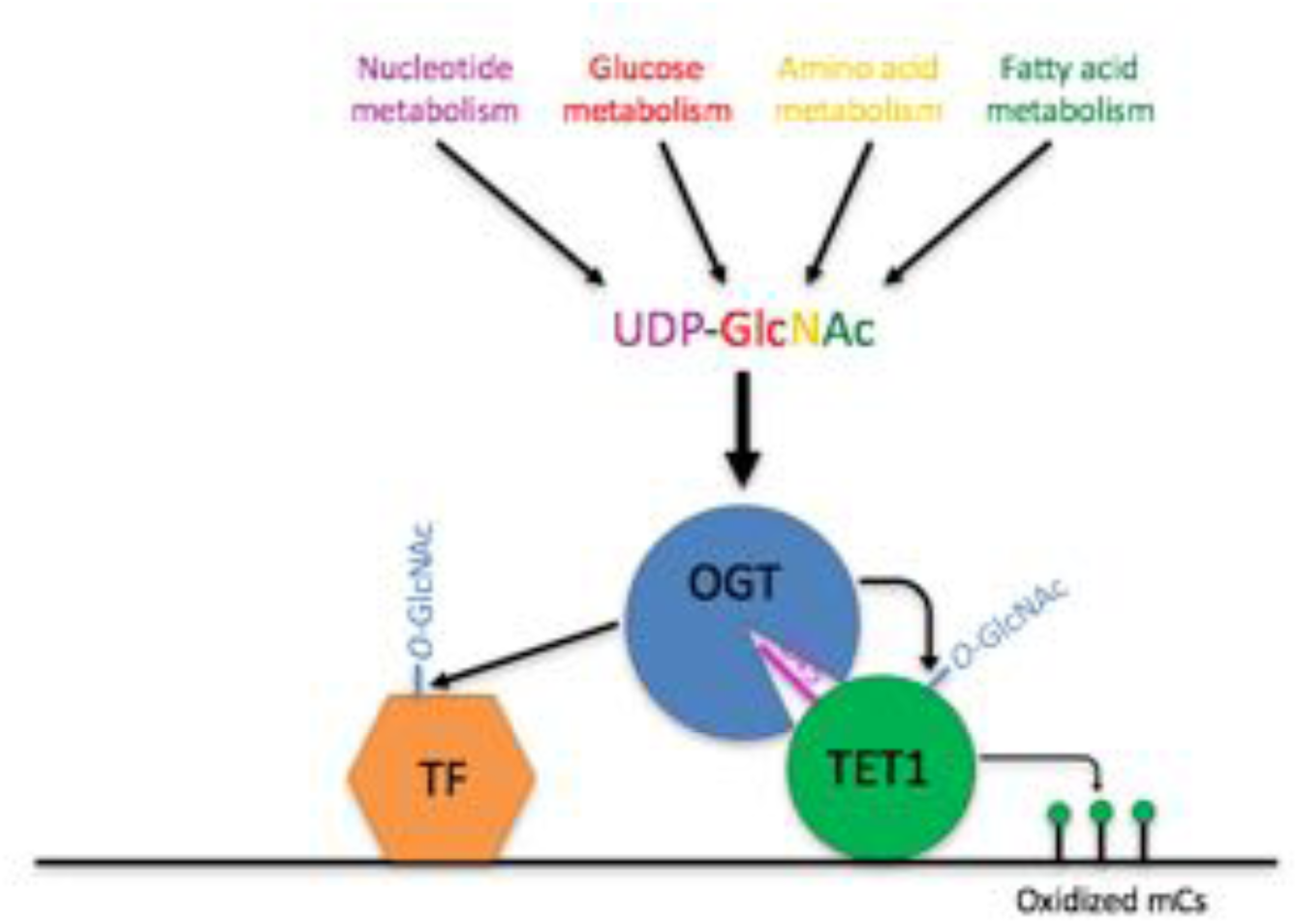
Model showing two roles of the TET1-OGT interaction in regulation of gene expression. OGT’s activity is regulated by the abundance of its cofactor UDP-GlcNAc, whose synthesis depends on the abundance of various cellular metabolites. OGT (blue circle) binds to TET1 (large green circle) via the TET1 C45 (purple line). OGT modifies TET1 and regulates its catalytic activity ( small green circles representing modified cytosines). At the same time, TET1 binding to DNA brings OGT into proximity of other DNA-bound transcription factors ( orange hexagon), which OGT also modifies and regulates.

The D2018A mutation reduced, but did not completely disrupt, the OGT-TET1 interaction in mESCs. In this cell type, TET1 is found in high molecular weight complexes that contain OGT, TET2, and the epigenetic regulator HCF1 [25]. In addition to TET1, OGT can directly interact with TET2 [26,27] and HCF1 [44] and TET2 co-IPs with TET1 [25]. Given the robust inhibition of the OGT-TET1 interaction by the D2018A mutation in multiple other contexts, our data are consistent with a scenario in which the D2018A mutation disrupts the direct interaction between OGT and TET1 in mESCs. However, some OGT remains associated in a co-IP assay through interactions with TET2 and/or HCF1. It is also possible that OGT interacts with the N-terminus of TET1, so that the D2018A mutation cripples but does not prevent the interaction with full-length TET1. This scenario is unlikely, however, since multiple studies have failed to identify any OGT binding capability outside the catalytic domains of TETs [27,34,45].

### Regulation of TETs by OGT in development

Our result that wild type TET1, but not TET1 carrying a mutation that can impair interaction with OGT, can rescue *tet2/3^DM^* zebrafish suggests that OGT regulation of TET enzymes may play a role in development. The importance of both TET proteins and OGT in development has been thoroughly established. Zebrafish lacking *tet2* and *tet3* die as larvae [30], and knockout of *Tet* genes in mice yields developmental phenotypes of varying severities, with knockout of all three *Tets* together being embryonic lethal [33,37,38,46]. Similarly, OGT is absolutely essential for development in mice [47], zebrafish [48], and *Drosophila* [49], though its vast number of targets have made it difficult to narrow down more specifically why OGT is necessary. Our results suggest that the function of the TET-OGT interaction in development may be two-fold. First, it may regulate TET enzyme activity. Second, OGT association with TETs may direct OGT to specific regions of the genome, facilitating *O*-GlcNAcylation of other transcriptional regulators in a site-specific manner (Fig. 7).

### A connection between metabolism and the epigenome

OGT has been proposed to act as a metabolic sensor because its cofactor, UDP-GlcNAc, is synthesized via the hexosamine biosynthetic pathway (HBP), which is fed by pathways metabolizing glucose, amino acids, fatty acids, and nucleotides [23]. UDP-GlcNAc levels change in response to flux through these pathways [50-52], leading to the hypothesis that OGT activity may vary in response to the nutrient status of the cell. Thus the enhancement of TET1 activity by OGT and the significant overlap of the two enzymes on chromatin [25] suggest a model in which OGT may regulate the epigenome in response to nutrient status by controlling TET1 activity (Fig. 7).

## Acknowledgments

We thank Diego Pasini for the OGT++ mESCs, Miguel Ramalho-Santos for the FLAG-TET1 CD plasmid, and Suzanne Walker for the His-OGT plasmid. We thank Leeanne Goodrich, Richard Yan, Christopher Agnew, and Sy Redding for technical assistance. We thank all members of the Panning lab for valuable ideas and discussion. This work was supported by R01 GM088506 (BP), NCI grant P30 CA008748 (MG), and funding from the Geoffrey Beene Cancer Research Center of Memorial Sloan-Kettering Cancer Center (MG). JH was supported by the California Institute for Regenerative Medicine Predoctoral Fellowship TG2-01153.

---

**Fig. S1:**
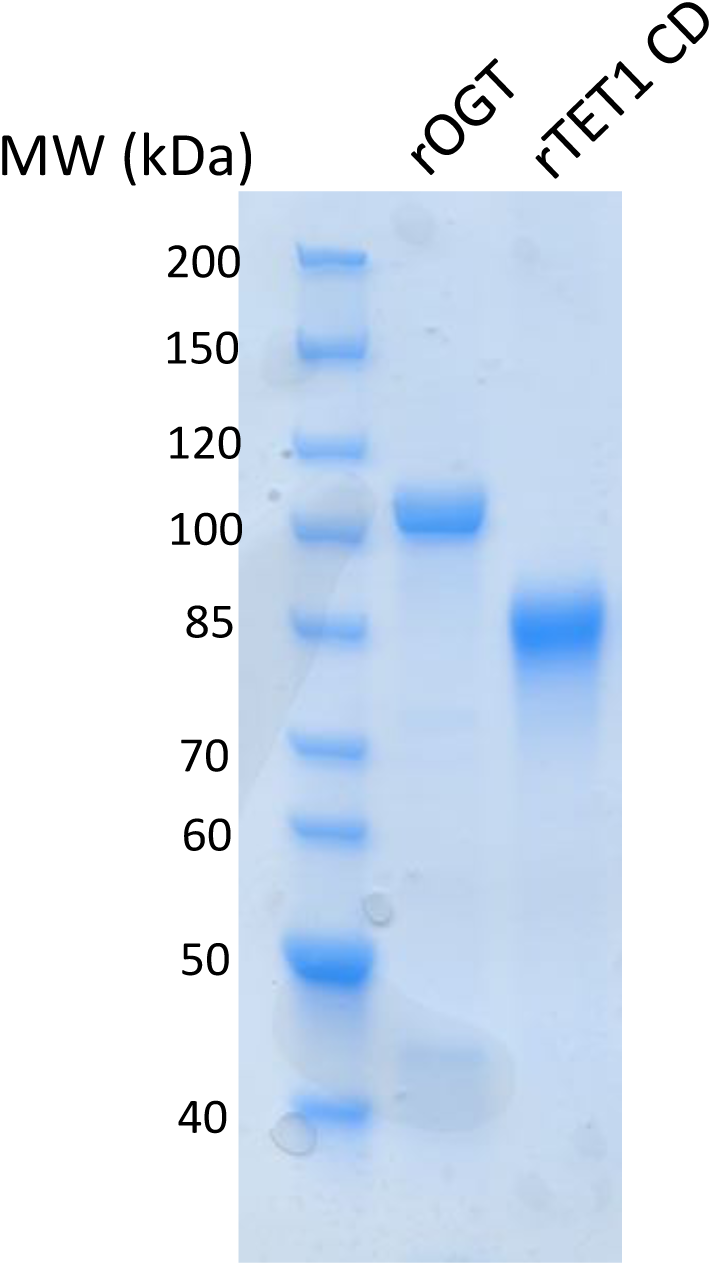
Coomassie stained SDS-PAGE gel of purified rOGT and rTET1 CD.

**Fig. S2:**
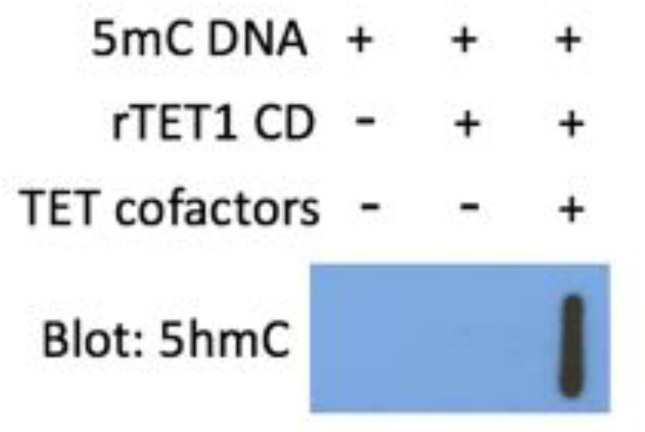
5hmC slot blot of biotinylated 5mC containing lambda DNA from rTET1 CD activity assays.

**Fig. S3:**
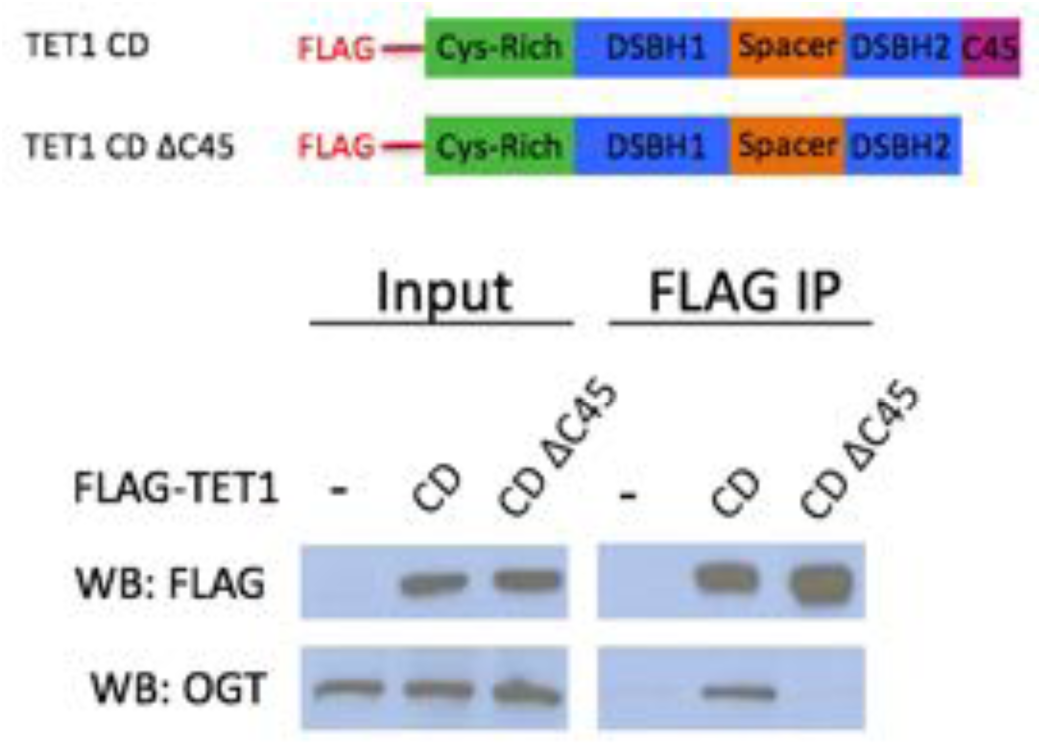
FLAG and OGT western blot of inputs and FLAG IPs from HEK293T cells transiently expressing FLAG-TET1 CD or FLAG-TET1 CD ΔC45 ( diagrammed in the upper panel).

**Fig. S4:**
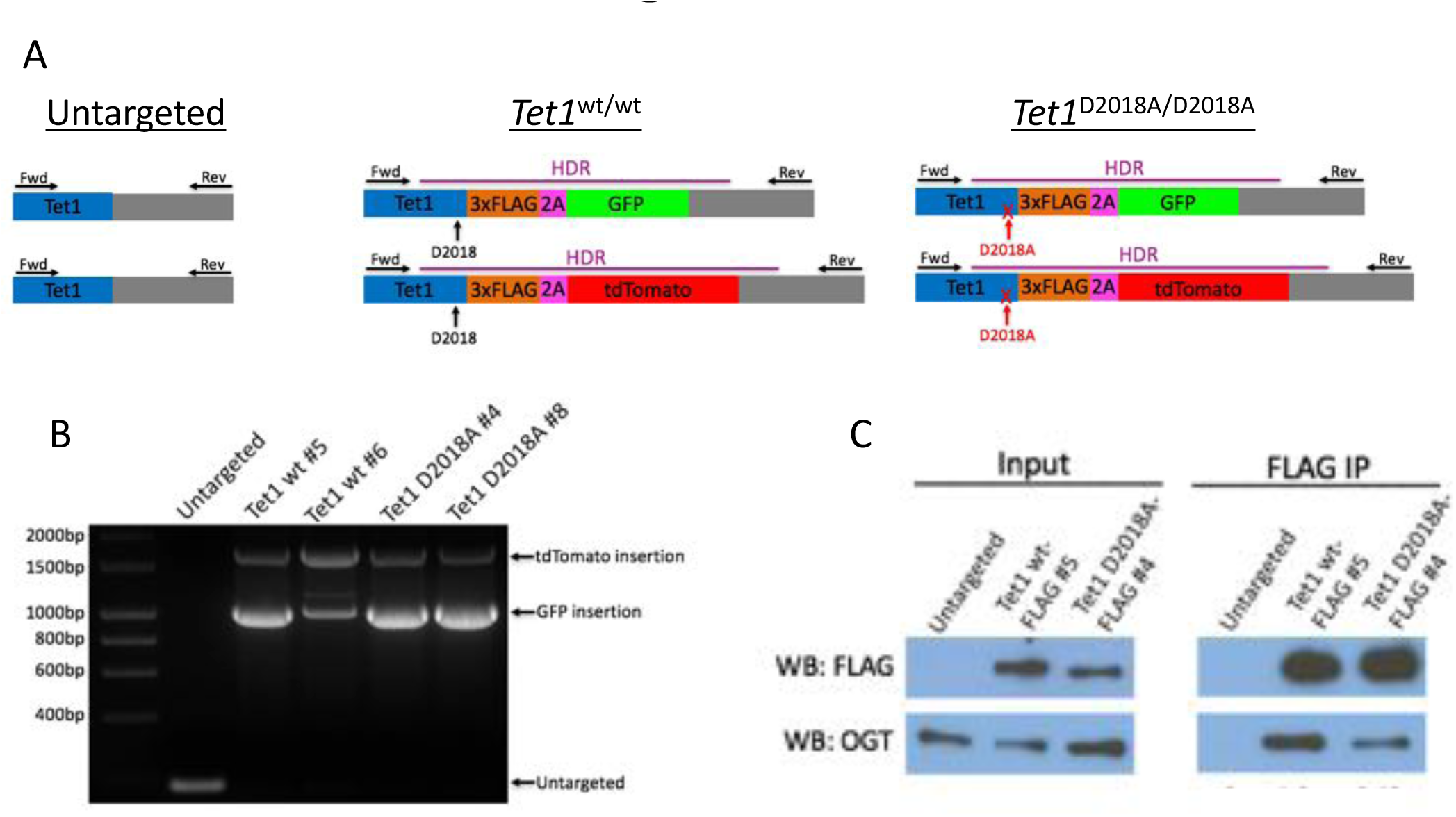
A) Schematic of mESC lines. DNA encoding a 3xFLAG tag was added to the 3’ end of both alleles of *Tet1*, followed by a 2A sequence and a fluorescent protein (GFP or tdTomato). The 2A sequence causes ribosome skipping, resulting in separate translation of TET1-3xFLAG and 2A-GFP or 2A-tdTomato. Purple line: template used for homology-directed repair (HDR). Horizontal arrows: primers used for PCR genotyping. Vertical arrows: D2018 residue. B) PCR genotyping of independently derived, clonal, targeted mESC lines using primers indicated in A. C) FLAG and OGT western blot of inputs and FLAG IPs from a second set of mESC lines in which a FLAG tag or a FLAG tag and the D2018A mutation were introduced into both copies of *Tet1*.

**Table S1:**
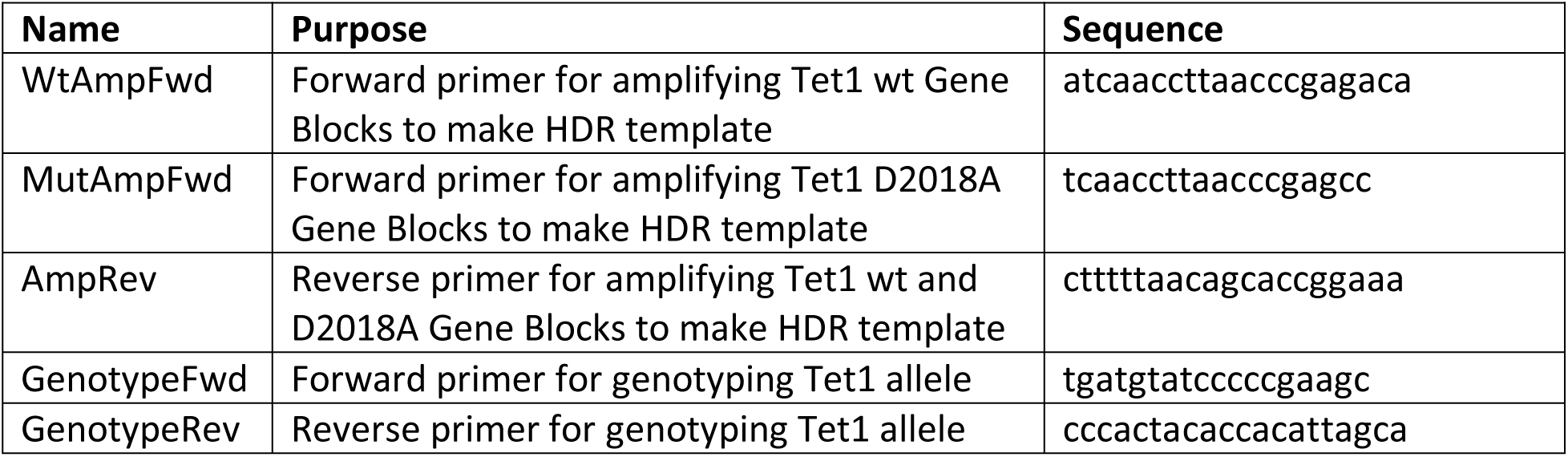
Primers used for creating and genotyping mESC lines.

**Table S2:**
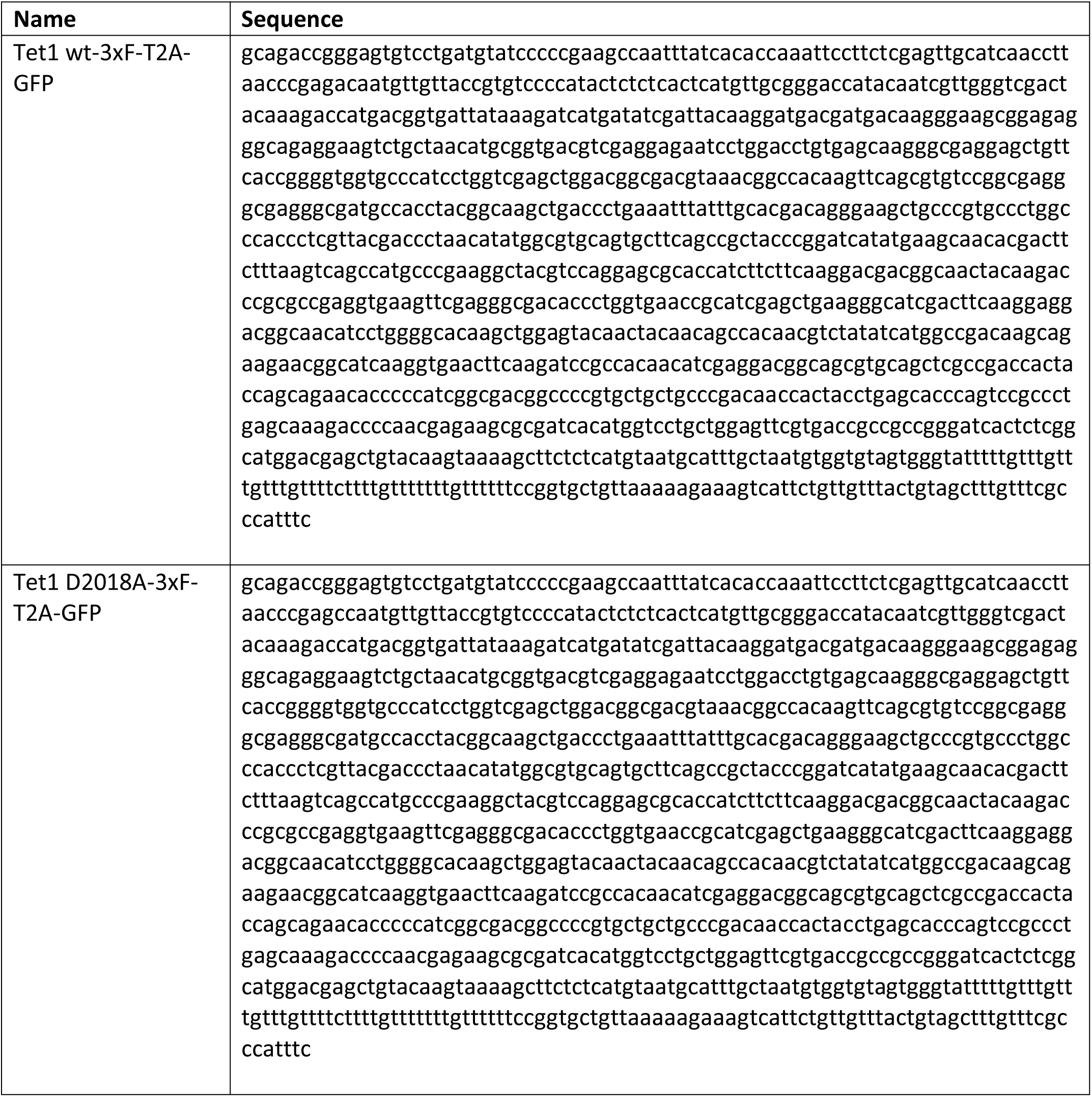
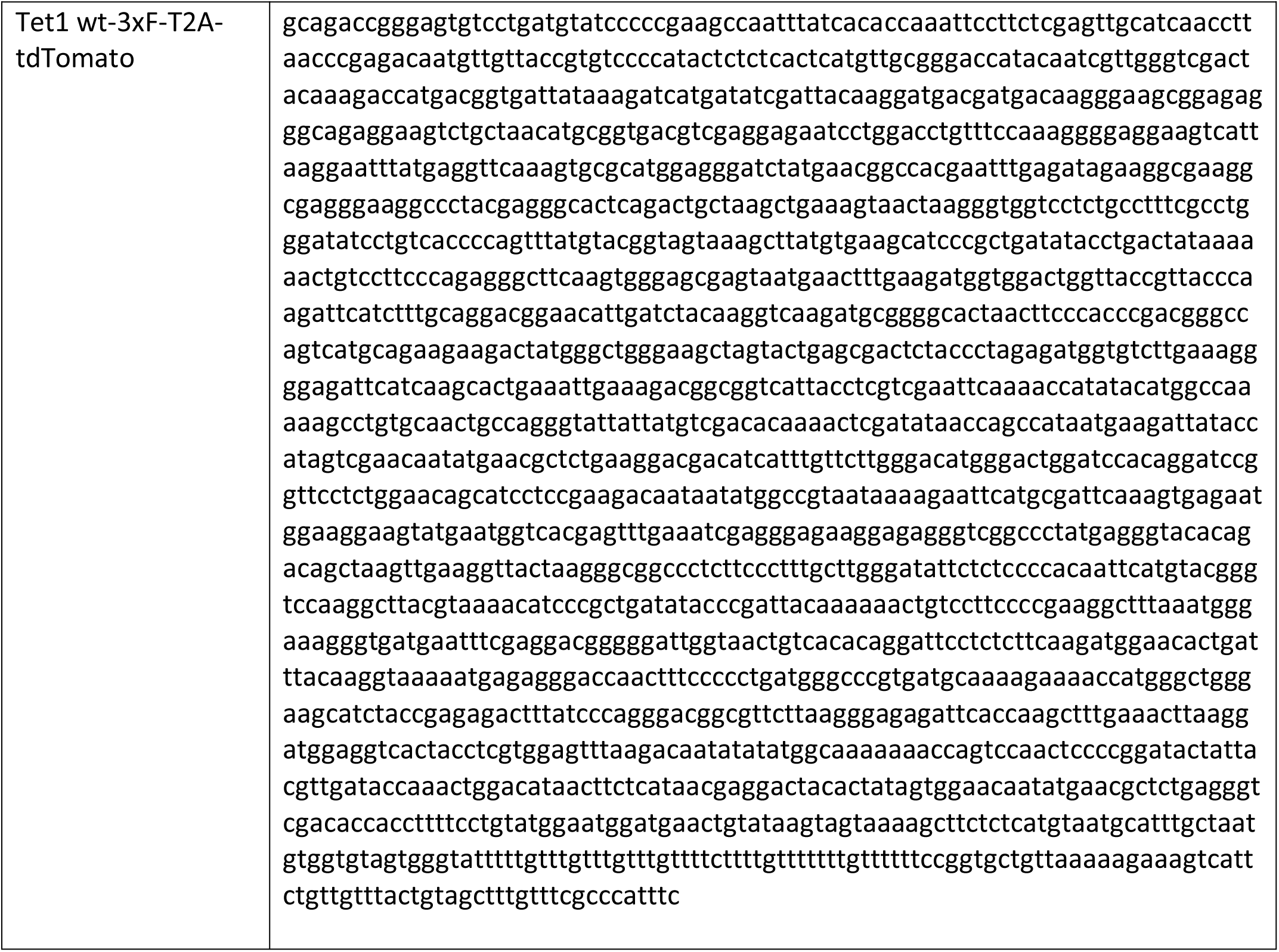
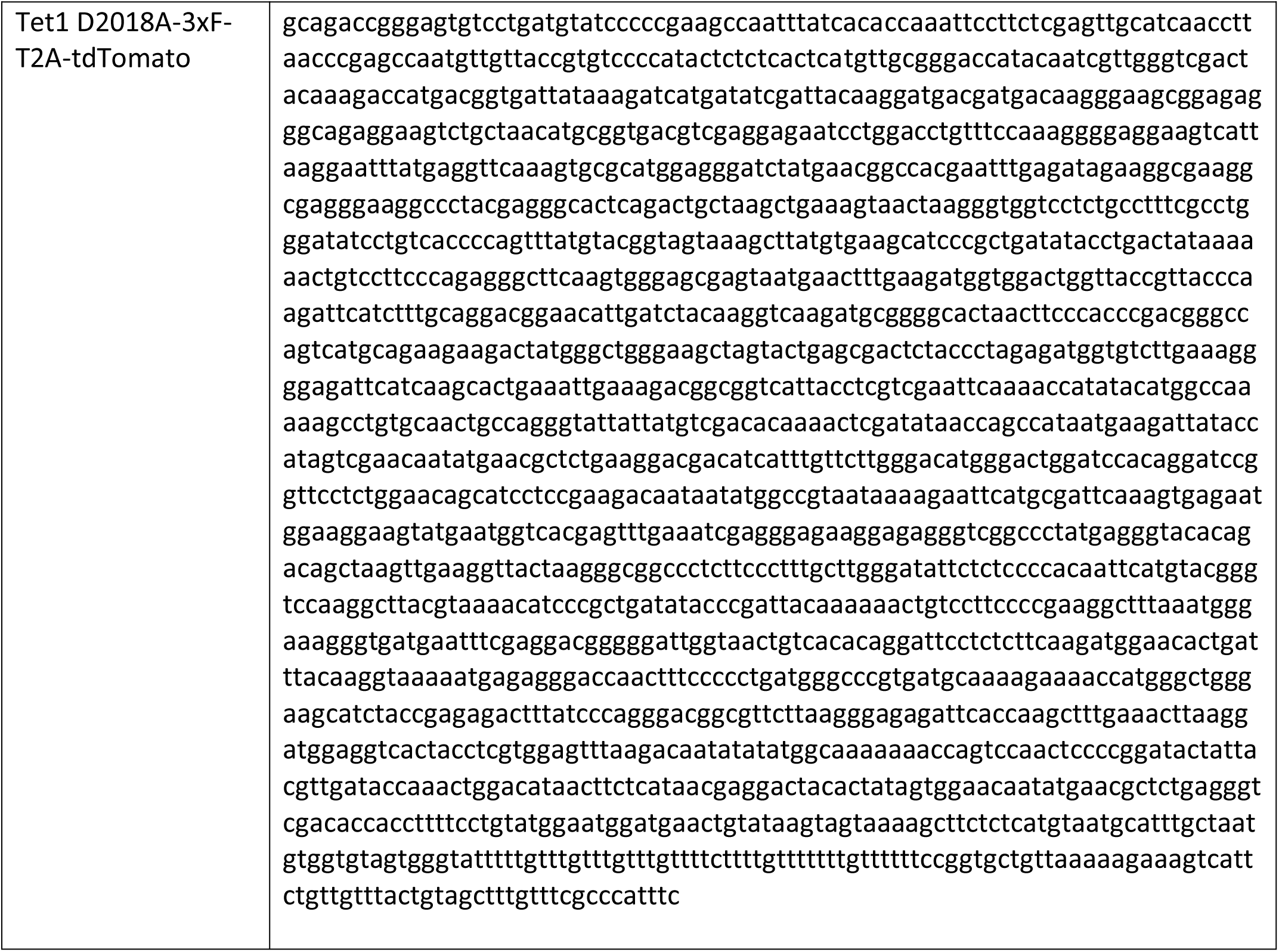
Gene blocks amplified to make HDR templates.

